# Numerical reproduction of the Sherrington-Adrian observations through a community of McCulloch-Pitts neurons with plastic remodelling

**DOI:** 10.1101/2023.12.05.570084

**Authors:** Luis Irastorza-Valera, José María Benítez, Francisco J. Montans, Luis Saucedo-Mora

**Affiliations:** E.T.S. de Ingeniería Aeronáutica y del Espacio, Universidad Politécnica de Madrid, Pza. Cardenal Cisneros 3, 28040, Madrid, Spain; Department of Materials, University of Oxford, Parks Road, Oxford, OX1 3PJ, UK; Department of Nuclear Science and Engineering, Massachusetts Institute of Technology, MA02139, USA; PIMM Lab, ENSAM (Arts et Métiers ParisTech, 151 Bd de l’Hôpital, Paris, France; Department of Mechanical and Aerospace Engineering, Herbert Wertheim College of Engineering, University of Florida, FL32611, USA

**Keywords:** neuronal model, plastic remodelling, signal homogenization, latency

## Abstract

Neurons form a highly complex network that produces cognition from simple associative rules. From previous results, this work shows the natural capability of the numerical network produced to modulate the output signal with independence of the intensity of the stimuli. Moreover, the plastic remodelling implemented in the model is capable to change the latency of a wide range of stimuli to synchronize them and adjust to a required delay of the signal.

## 1. Methodology and results

Regarding brain activity, Adrian observed that with independence of the amplitude of stimuli, the amplitude of the response was unaffected [2]. Latter observations demonstrated that the impedance of the brain is independent of the frequency of the stimulation [9]. Other works, such as McNamara et al. [11] shows this tendency of the neuronal circuits to stabilize different input signals, and its implication in diseases such as Parkinson. This stability also have implications in disorders generated by Glioblastoma multiforme [5], which generates a variation of the mechanical loads in certain regions of the brain.

This tendency of the brain has been modelled [12] [3] to reproduce the stabilization of signals through noise inserted in the neuronal circuit. And supported by experimental results [4], concluded that there is an adaptation in the neuronal network that tends to stabilize the response of the system. Therefore, the concept of metastability [13] is aligned with those results. For the Sherrington-Adrian observations [2] [8], the authors received the nobel prize of medicine in 1932 arguing that they “discovered that the explosive waves of impulses discharged along the nerve are always the same size, regardless of how strong the stimulus is” [1].

The model proposed in this work is a neuronal network based on simple McCulloch-Pitts neurons [10], which are capable to do plastic remodelling to adapt to a required signal delay [6]. The model is a sphere with a radius of 2 mm, and 40.000 neurons, which is lower than the neuronal density of the brain that can be considered as 40.000 neurons/*mm*^3^ [7]. The sphere has 1.000 neurons in the top where the excitation begins, and 1.000 neurons on the bottom in charge of the supervision of the plastic remodelling [6]. Figure 1 shows the results of the model when different signals within a full range of intensity are used to excite the same neuronal network.

**Figure 1.**
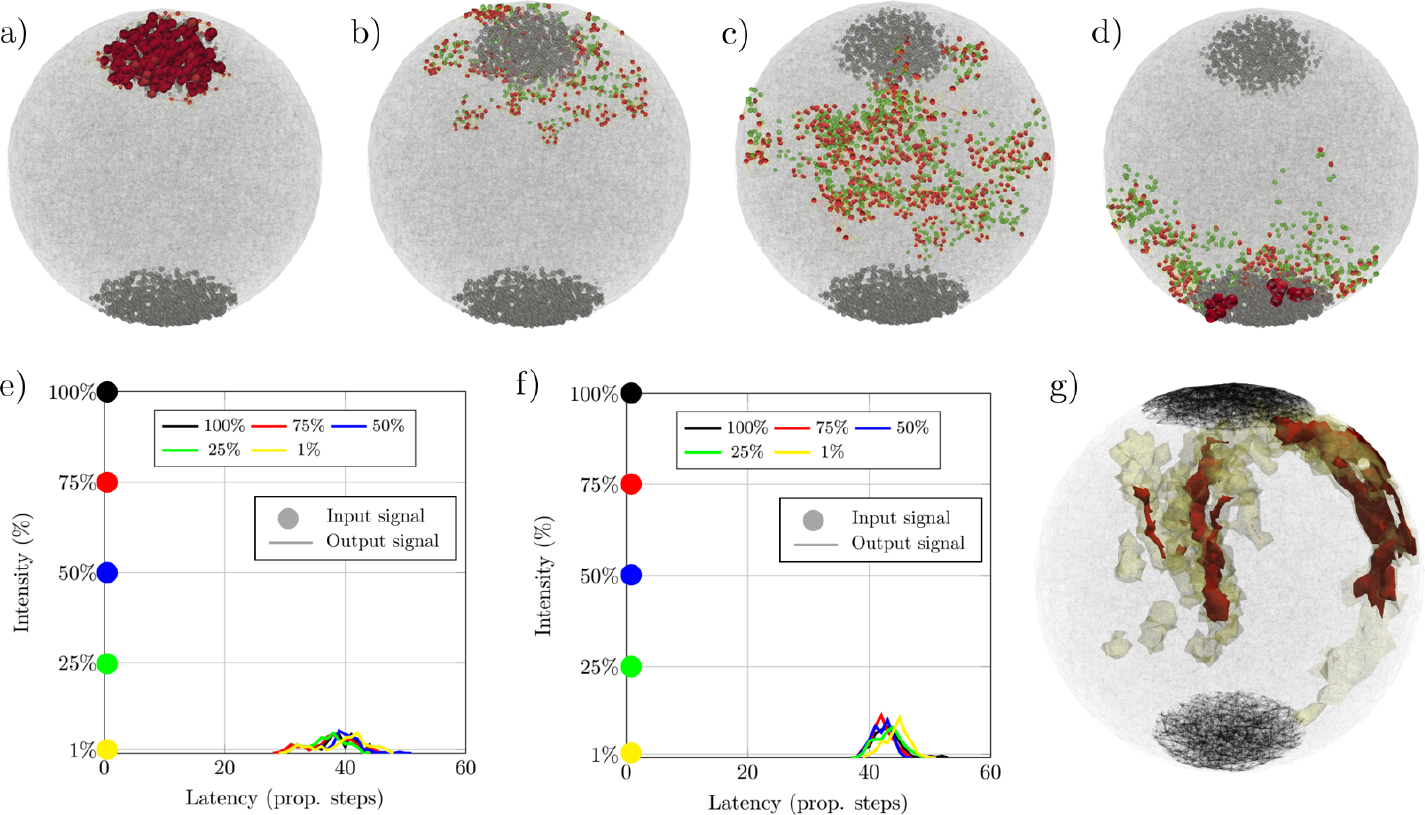
Signal propagation through a network of McCulloch-Pitts neurons, and its plastic remodelling. From a) to d) we show the propagation of a signal with the 100 % of initial stimulation, and remodelled, at the steps of propagation 0 (a), 5 (b), 15 (c) and 35 (d), the red dots are the neurons fired at this step, and the green dots their pre-synaptic neurons. The remodelled peak of the signals is at 45 steps of propagation. e) shows the original signal outputs of the network under different stimuli, and f) are the remodelled outputs of the network, which are intended to be the same and centred at 45 steps of propagation. Those output curves are the density of output neurons that are excited at each step. g) shows the regions where the remodelled neurons are, in this case, for an early global plastic remodelling. Here in g) the red region shows the zone with higher density of remodelling, and the yellow one has a lower amount. If we consider the propagation speed of the signal as 10 *m/s*, each step of propagation of this figure can be assumed as around 5 *μs*.

The preliminary result of Figure 1 shows the capability of the network to modulate the signals and to produce the same output excitation regardless of the intensity of the stimuli, which ranges from 100% of the neurons excited (i.e. 1.000) to a 1% (i.e. 10 neurons). Figure 1a-d shows the case with the 100% of the input neurons excited, and the colors denote the neurons fired (red) and the pre-synaptic neurons of those (green). In all the cases, with different stimuli, the output pulse is equal in size and latency.

## 2. Conclusions

The presented work shows the natural capability of this model to modulate signals with input stimulus of different intensity, and that it is done even with a low number of McCulloch-Pitts neurons. Also, we demonstrate that through plastic remodelling the neuronal network can synchronize and modulate the responses to different stimuli, and even simultaneously adapt them to a required signal delay.

## Declarations

### Ethics approval and consent to participate

Not applicable.

### Consent for publication

Not applicable.

### Availability of data and materials

The data will be provided under request to the corresponding author.

## Funding

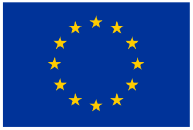

*This project has received funding from the European Union’s Horizon 2020 research and innovation programme under the Marie Slodowska-Curie Grant Agreement No. 956401*

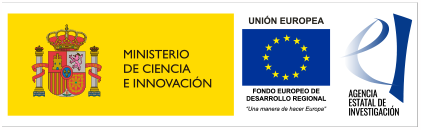

*Grant PID2021-126051OB-C43 funded by MCIN/AEI/ 10*.*13039/501100011033 and by “ERDF A way of making Europe”*.

## Authors’ contributions

LIV crated the new software used in the work and drafted the work. JMB interpretation of data and revision. FJM interpretation of data and revision. LSM conception and design of the work, creation of new software used in the work, interpretation of data and revision. All authors approved the final Manuscript

## Acknowledgements

Not applicable.

